# Intestinal mucus and gut-vascular barrier: FxR-modulated entry sites for pathological bacterial translocation in liver cirrhosis

**DOI:** 10.1101/690057

**Authors:** M. Sorribas, M. O. Jakob, B. Yilmaz, H. Li, D. Stutz, Y. Noser, A. de Gottardi, S. Moghadamrad, M. Hassan, A. Albillos, R. Francés, O. Juanola, I. Spadoni, M. Rescigno, R. Wiest

**Affiliations:** Maurice Müller Laboratories, Department for Biomedical Research, University of Bern, Bern, Switzerland; Department of Visceral Surgery and Medicine, Bern University Hospital, University of Bern, Bern, Switzerland; Department of Gastroenterology, Hospital Universitario Ramón y Cajal, IRYCIS, Madrid, Spain; Hepatic and Intestinal Immunobiology Group, Universidad Miguel Hernández-CIBERehd, San Juan, Spain; Humanitas University, Department of Biomedical Sciences, Via Rita Levi Montalcini, 20090 Pieve Emanuele, MI, Italy; Humanitas Clinical and Research Center, Via Manzoni 56, 20089 Rozzano, MI, Italy

**Keywords:** portal hypertension, liver cirrhosis, gut-liver-axis, gut-vascular barrier, intestinal permeability, mucus, FXR

## Abstract

**Background and aims:** Pathological bacterial translocation (PBT) in liver cirrhosis (LC) is the hallmark for spontaneous bacterial infections increasing mortality several-fold. Factors known to contribute to PBT in LC are among others an increased intestinal permeability of which however, the mucus layer has not been addressed so far in detail. A clear route of translocation for luminal intestinal bacteria is yet to be defined but we hypothesize that the recently described gut vascular barrier (GVB) is impaired in experimental portal hypertension leading to increased accessibility of the vascular compartment for translocating bacteria.

**Results:** Healthy and pre-hepatic portal-hypertensive (PPVL) mice lack translocation of FITC-dextran and GFP-*Escherichia coli* from the small intestine to the liver whereas bile-duct-ligated (BDL) and CCl4-induced cirrhotic mice demonstrate pathological translocation which is not altered by prior thoracic-duct ligation. Mucus layer is reduced in thickness with loss of goblet-cells and Muc2-staining and expression in cirrhotic but not PPVL-mice associated with bacterial overgrowth in inner mucus layer and pathological translocation of GFP-*E.coli* through the ileal epithelium. GVB is profoundly altered in BDL and CCl4-mice with Ileal extravasation of large-sized 150 kDa-FITC-dextran but only minor in PPVL-mice. This pathological endothelial permeability and accessibility in cirrhotic mice associates with an augmented expression of PV1 in intestinal vessels. OCA but not fexaramine stabilizes the GVB whereas both FXR-agonists ameliorate gut-liver-translocation of GFP-*E.coli*.

**Conclusions:** Liver cirrhosis but not portal hypertension per se grossly impairs the endothelial and muco-epithelial barriers promoting PBT to the portal-venous circulation. Both barriers appear FXR-modulated with –agonists reducing PBT via the portal-venous route.

## Introduction

The gut-liver-axis presents the pathophysiological hallmark for initiation and/or perpetuation of multiple liver diseases^1^ and has been proposed to be fueled by pathological bacterial translocation (PBT) from the gut^2^. In liver cirrhosis, PBT from the gut into the liver and systemic circulation is one of the causes of bacterial infections and the augmented pro-inflammatory response to gut-derived products^2, 3^. In fact, failure to control invading bacteria and bacterial products in concert with host susceptibility determines remote organ injury in liver cirrhosis. This may include acute-on-chronic liver failure, hepato-renal-syndrome and hepatic encephalopathy worsening prognosis^4^. PBT in liver cirrhosis has been attributed so far to small intestinal bacterial overgrowth, increased intestinal permeability and lack of host defense mechanisms^5^. Here we focused on the first and last barrier separating luminal bacteria and vascular compartment namely, intestinal mucus and the newly defined gut-vascular barrier (GVB)^6^, both not having been addressed so far in liver cirrhosis and PBT.

Mucus represents the first frontier that commensal microbes in the gut have to cross in order to achieve PBT. The mucus consists of two layers with a similar protein composition where mucin-2 (MUC2) is the main component^7^. On one hand, the inner mucus layer is firmly attached to the epithelium, is densely packed and is devoid of bacteria^8^. On the other hand, the outer mucus layer is much more mobile, looser and is colonized with a distinct bacterial community^9^. Goblet cells are responsible for the formation of both the inner and outer mucus layer^10^ but also sense bacteria^11^ and react accordingly with mucin secretion^12, 13^. After crossing the mucus and epithelial barrier, translocating bacteria reach the lymphatic system as shown by culture positive mesenteric lymph nodes in experimental cirrhosis in multiple independent studies^14, 15^. In contrast, access to the intestinal microcirculation and portal-venous route has been proposed for PBT^16^, but has not been delineated in detail in portal hypertension and liver cirrhosis yet. The splanchnic circulation in portal hypertension presents with multiple vascular abnormalities^17^ including arterial vasodilation^14^, hyporesponsiveness to vasoconstrictors^18, 19^ and increased angiogenesis^20^. However, accessability of the intestinal microcirculation and thus, portal-venous route has not been investigated in portal hypertension so far. Endothelial barriers are characterized by the presence of junctional complexes which strictly control paracellular trafficking of solutes, fluids and cells^21^. In healthy conditions, the endothelial vascular barrier discriminates between differently sized particles of the same nature with 4kDa-dextran freely diffusing through the endothelium, whereas 70kDa-dextran does not. Plasmalemma-vesicle-associated protein (PV)-1 is an endothelial cell-specific protein that forms the stomatal and fenestral diaphragms of blood vessels^22^ and regulates basal permeability ^23^.

Liver cirrhosis is characterized by deficient levels of luminal bile acids in the gut^24^. Bile acids have been long known for their major effects on the microbiome and the intestinal barrier function. They exert their effects via transcription factors among which the Farnesoid X receptor (FXR) known to be one of the most important. FXR-activation has been reported to influence epithelial cell proliferation^25^ and to exert potent anti-inflammatory actions in the intestine, stabilizing epithelial integrity^26–29^. Moreover, FXR stimulation in the small intestine exerts antibacterial actions via induction of antimicrobial substances^30^ and FXR-agonists have been shown to ameliorate chemically induced intestinal inflammation with improvement of colitis symptoms and inhibition of epithelial permeability. However, the exact role of bile acids and FXR in controlling intestinal muco-epithelial as well as vascular permeability is still unknown. In addition, the microbiome has been proposed to play a key role for mucus-synthesis, release and barrier-function^31^ but information on its impact on goblet cell density and mucus thickness are limited. Finally, although colonization by microbial commensals is known to promote vascular development^32^ its impact and modulatory role on the GVB-function is not known.

Taken together, the aims of the current study were i) to characterize changes in mucus barrier as well as GVB in germ-free conditions and in the context of liver cirrhosis or portal hypertension as well as ii) to delineate PBT from the gut to the liver along the gut-liver-axis in liver cirrhosis and iii) to unravel the role(s) of FXR for PBT, mucus- and gut-vascular-barrier.

## Methods

### Mice and animal models

Female C57BL/6J mice were purchased from ENVIGO (Horst, The Netherlands) and kept at the Central Animal Facility of the University of Bern under specific pathogen-free conditions. <u>Mice were kept in next-generation IVC cages with enriched environment, 12 hour day-night cycle and fed ad libitum. All</u> experiments involving animals were performed in accordance with Swiss Federal regulations and with local institutional approval. Where indicated, animals were kept at either germ-free conditions or at gnotobiotic conditions with a stable defined moderately diverse mouse microbiota (sDMDM2) that consists of 12 bacterial species which has been developed and used previously in our lab^9^. The intestine-specific FXR-null (FXR⊗^IE^) mice are on a C57BL/6 genetic background and kindly provided by Prof. B. Schnabl from University of California San Diego, USA.

***Bile-duct-ligation***: Bile Duct Ligation (BDL) was performed as previously described^33^. Briefly, in an aseptic setting the abdomen was incised in the midline under isoflurane anesthesia and administration of buprenorphine 60/µg/kg body weight. The bile duct then cleared from adjacent structures. Three different ligatures were placed along the bile duct with a non-absorbable suture and the bile duct was resected between the second and third ligature. Sham-operated mice were treated similarly but no ligature was placed. Experiments were performed 10 days to 4 weeks after surgery. ***Partial-Portal-Vein Ligation***: To induce pre-hepatic portal hypertension, partial portal vein ligation (PPVL) was performed as follows: In an aseptic setting the abdomen was incised in the midline under isoflurane anesthesia and administration of buprenorphine 60/µg/kg body weight. Portal vein was cleared from surrounding tissue. To achieve only partial and not complete ligation, a non-absorbable suture was placed around a 26-gauge blunt-tipped needle lying alongside the portal vein. After ligation the blunt-tipped needle was removed, leaving a calibrated stenosis on the portal vein. Sham-operated mice were treated the same with no ligature placed. Experiments were performed 14 days after surgery. ***CCl4-induced cirrhosis***: Cirrhosis was induced according to the method described before in rats (Runyon, Sugano, Kanel, & Mellencamp, 1991) and modified for mice^34^. After 1 week of phenobarbital administration in the drinking water (0.3g/L), CCl4 inhalation was started: Mice were placed three times per week in a gas chamber and compressed air, bubbling through a flask containing CCl4, was passed into the gas chamber via a flowmeter (2 L/min). Afterwards, bubbling was stopped and animals were let breath normal air. The length of CCl4 exposure was progressively increased, 1 min of bubbling the first week, 1.5 min of bubbling in the second week, and then 2 min of bubbling throughout all the rest of the experiment. Mice were treated with 1 cycle/treatment in the initial 3 weeks, with 2 cycles/treatment in the fourth week, and with 3 cycles/treatment thereafter. Once treatment consisted of two or more cycles, these were separated by at least 10 minutes of breathing in ambient air not containing CCl4. ***Thoracic duct ligation***: Thoracic duct ligation was performed as previously described with some alterations^35^. After gavage of an oil solution 30 minutes prior to the operation, mice were anesthesized and a left-sided subcostal incision was performed. After incision of the dorsal retroperitoneum, the aorta was freed from surrounding fatty tissue. The thoracic duct was exposed and carefully freed from the aorta. A metal clip was then placed for closure of the thoracic duct (Clip 9 Vitalitec, Peters Surgical, Bobigny Cedex, France). The incision was closed with a two-layer running suture (Prolene 6-0, Ethicon, Johnson & Johnson, Spreitenbach, Switzerland).

*FxR-modulation pharmacologically* was achieved by high-affinity FXR-agonists, namely fexaramine (Fex)^36^ and obeticholic acid (OCA)^37^ treatment by oral gavage. Fex is known to lack major absorption and systemic appearance (< 30nm after per oral 100 mg/kg/day) and thus is rendered intestinal-specific meaning luminally and hence active at the muco-epithelial site^38^. In contrast, OCA per os is rapidly and highly absorbed achieving sufficient systemic drug levels (> 10 µmol in serum after 30 mg/kg/day^39^) thus targeting muco-epithelial as well as endothelial and hence, gut-vascular barrier.

### In-vivo permeability and intestinal loop assay

Mice were placed under isoflurane anesthesia middle line laparotomy was performed and terminal ileum exteriorized and 1cm long of terminal ileum ligated. 8mg of 4 KDa FITC-Dextran (Sigma-Aldrich) or GFP-E.coli (10^5^ CFU) were injected in the loop dissolved in 200uL of saline. 1 hour after injection liver, mesenteric lymph nodes (mLN), spleen and the intestinal loop were harvested and fixed in Carnoy’s fixative (60% Ethanol, 30% Chloroform, 10% glacial acetic acid) and embedded in paraffin. The amount of the dye was measured in the liver, spleen and mLN by immunofluorescence of 5um sections stained with 4’,6-diamidin-2-fenilindolo (DAPI). Number of particles (FITC-signal or GFP-*E.coli*) were counted per high-power field at 40X in a fluorescence microscope (Nikon, Eclipse Ti-E). In some experiments, prior to isolation of the loop and inoculation of FITC-dextran or GFP-E.*coli* the thoracic duct was ligated to avoid any lymphatic drainage of translocating agents into the systemic circulation that otherwise could contribute to the hepatic counts. To control for tissue auto-fluorescence some intestinal loops were injected with saline and no particles were count in the tissue harvested from these mice.

### In-vivo intravital probe-based confocal laser endomicroscopy

Mice were placed under isoflurane anesthesia 1 cm long incision was made on the skin to expose the peritoneal wall. An additional small incision was made on the peritoneum to expose the abdominal cavity. A 3- to 4-cm loop from the ileum was externalized and endomicroscopy probe (ColoFlex™ UHD, Mauna Kea Technologies) was fixed at an angle of 60° from the operation table close to the exposed intestinal loop. A small incision was made in the anti-mesenteric surface of the intestinal wall and the endomicroscopy probe was inserted in the intestine. The probe was placed so that at least one villus was visualized on the screen. After stable imaging was achieved a retro-orbital intravenous injection was made with a FITC-Dextran solution (50µL of 40mg/mL for 4kDa-FITC-Dextran, 50µL of 10mg/mL for 40 kDa and 70 kDa-FITC-Dextran or 100µL of 10mg/mL for 150kDa-FITC-Dextran solution from SIGMA). Immediately after the injection the FITC signal was observed in the intestinal villi capillaries and the intestine was pulled using forceps in order to scan the epithelial surface. Endoscopy video was recorded for 30 minutes of total observation time.

### Endothelial leakage analysis from *in vivo* endomicroscopy

Cellvizio Viewer software (Mauna Kea Technologies) was used in order to extract the photograms from the endomicroscopy video. The images with focused vessels and normal blood flow were selected. At least one picture per time point was selected and 7 measurements were taken per picture. Measurements were made using Image J as follows: straight lines perpendicular to the vessel were drawn in each vessel within the picture and grey value measured in each pixel along the line. Mean grey value (MGV) of the vessel was calculated and background fluorescence was extracted, MGV of the immediate extravascular space was calculated in the same way. To avoid a bias in our measurements we took the same numbers of pixels analyzed from the intra and extra-vascular space in all the mice. We also avoid measuring these extravascular spaces where more than one vessel could contribute to the fluorescence values. Results are shown as ratio between mean of extravascular fluorescence and mean of intravascular fluorescence. When extravasation was too high to distinguish vessels from lamina propria a value of 1 was given arbitrary.

### Inter-epithelial leakage analysis from *in vivo* endomicroscopy

Cellvizio Viewer software (Mauna Kea Technologies) was used in order to extract the photograms from the endomicroscopy video. Only these photograms where the vessels were in focus and properly perfused were selected. The first time when the majority of the interepithelial spaces were seen positive for fluorescein was noted. When this requirement was not accomplished a value of 30 minutes was given.

### Dual band confocal laser endomicroscopy in-vivo in intestinal loop experiments

Mice were anesthetized with an isofluorane-oxygen mixture. Laparotomy was performed and terminal ileum intestinal loop exposed. Endomicroscopy probe (UltraMiniO) was fixed at an angle of 60° from the operation table close to the exposed intestinal loop. A small incision was made in the anti-mesenteric surface of the intestinal wall and the endomicroscopy probe was inserted in the intestine. The probe was placed so that at least one villus was visualized on the screen. After stable imaging was achieved one ligature was placed 1cm proximal to the gut opening and another around the probe and the gut itself. After ligation retro-orbital intravenous injection of 100µL 2% evans blue (SIGMA) solution was performed. GFP-E.coli was injected to the loop (10^5^ CFU in 0.2 ml media). Visualization of *E.coli* and intestinal villi capillaries was performed using Cellvizio ® DualBand (Mauna Kea Technologies) for 1 hour.

### Immunofluorescence and confocal microscopy for endothelial cell-to-cell-interactions and plasmalemma vesicle-associated protein-1 (PV1)

Intestinal samples were fixed overnight in paraformaldehyde, L-Lysine pH 7.4 and NaIO4 (PLP buffer). They were then washed, dehydrated in 20% sucrose overnight and included in OCT compound. 10um cryosections were rehydrated, blocked with 0.1M Tris-HCl pH 7.4, 2% FBS, 0.3% Triton X-100 and stained with the following antibodies: anti-mouse PLVAP (clone MECA32, BD Pharmingen), anti-ZO1 (clone ZO1-1A12, Invitrogen), anti-Occludin (clone OC-3F10, Invitrogen), anti-mouse CD34 (clone RAM34, eBioscience). Slices were then incubated with the appropriate fluorophore-conjugated secondary antibody. Before imaging, nuclei were counterstained with DAPI. Confocal microscopy was performed on a Leica TCS SP5 laser confocal scanner mounted on a Leica DMI 6000B inverted microscope equipped with motorized stage. Image J software package was used for image analysis and fluorescence quantification.

### Goblet cell (GC) analysis

1cm long sections of different parts of the small intestine (duodenum, jejunum and ileum) and colon were fixed in Carnoy’s fixative (60% Methanol, 30% Chloroform and 10% Glacial acetic acid) overnight and embedded in paraffin. 6µm thin cuts were deparaffinized and GC stained with periodic acid-Schiff solution (PAS) and hematoxylin to stain nuclei. Pictures were taken at 200X magnification with a bright field microscope GC were counted and villi perimeter measured using Image J. We have demonstrated that perimeter of the villi strongly correlates (R^2^=0.9028) with the number of epithelial cells (data not shown). Hence the ratio between number of GC and perimeter is representative of the proportion of goblet cells amongst epithelial cells in a given villus.

### Mucus parameters

Mucus thickness measurements in gut explants were performed using a micromanipulator holding a cell injection pipette with the corresponding glass capillary tilted to a delivered angle. The position of the capillary was recorded when touching the mucus layer and recorded again when touching the villus, visualized under a stereomicroscope. Mucus thickness was calculated via trigonometrical approach as the product between the measured distance and the sinus of the given angle. 5 measurements per organ were performed. Inner mucus layer in sDMDM2-conditions was assessed for bacterial load after removing outer layer by suction being harvested manually and cultured on LB agar plates for 12 hours. Results were shown as colony forming unites per mg mucus.

### Electronmicroscopy

Mouse intestine was fixed with 2.5% glutaraldehyde (Agar Scientific, Stansted, Essex, UK) and 2% paraformaldehyde (Merck, Darmstadt, Germany) in 0.1M Na-cacodylate-buffer (Merck, Darmstadt, Germany) with a pH of 7.33. Samples were fixed for at least 24 hours before being further processed. Samples were then washed with 0.1M Na-cacodylate-buffer three times for 5 min, postfixed with 1% OsO4 (Electron Microscopy Sciences, Hatfield, USA) in 0.1 M Na-cacodylate-buffer at 4°C for 2 hours, and then washed in 0.05M maleic acid (Merck, Darmstadt, Germany) for three times 5 min.. Thereafter samples were dehydrated in in 70, 80, and 96% ethanol (Alcosuisse, Switzerland) for 15 min each at room temperature. Subsequently, the tissue was immersed in 100% ethanol (Merck, Darmstadt, Germany) for three times 10 min, in acetone (Merck, Darmstadt, Germany) for two times 10 min, and finally in acetone-Epon (1:1) overnight at room temperature. The next day, samples were embedded in Epon (Sigma-Aldrich, Buchs, Switzerland) and left to harden at 60°C for 5 days. Sections were produced with an ultramicrotome UC6 (Leica Microsystems, Vienna, Austria), first semithin sections (1um) for light microscopy witch were stained with a solution of 0.5% toluidine blue O (Merck, Darmstadt, Germany) and then ultrathin sections (75nm) for electron microscopy. The sections, mounted on 200mesh copper grids, were stained with uranyless (Electron Microscopy Sciences, Hatfield, USA) and lead citrate (Leica Microsystems, Vienna, Austria) with an ultrostainer (Leica Microsystems, Vienna, Austria). Sections were then examined with a transmission electron microscope (CM12, Philips, Eindhoven) equipped with a digital camera (Morada, Soft Imaging System, Münster, Germany) and image analysis software (iTEM).

### RNA Sequencing and network analysis

Whole tissue RNA was extracted from frozen ileum sections by homogenizing with Tissue Lyzer (QIAGEN) and TRIzol reagent (Life Technologies). 200uL of chloroform was added, samples mixed and centrifuged for. The upper RNA-containing phase was recovered and RNA precipitated with ice-cold isopropanol. The solution was centrifuged and RNA pellet washed with 75% ethanol. Once dried it was resuspended in RNase-free water. Concentration and purity was measured with Bioanalyzer (AGILENT). Library preparation was done using the RIboMinus protocol. Libraries were sequenced by Illumina HiSeq 3000 on the 100bp paired-end mode. The quality of the reads was documented as well as the number of them and they were mapped to the mouse genome. Number of reads per gene were compared to the different experimental groups using DESeq2 package from R software. Adjusted p-value and log2 fold change was calculated according to Benjamini and Hochberg, 1995. Protein-protein interaction was performed using STRIG database for these proteins which genes were significantly regulated (log2 fold change higher than 2 and adjusted p-value lower than 0.01).

## Statistical analysis

Statistical differences were evaluated using GraphPad Prism software Version7.0. The values were compared using either Student’s t-test or ANOVA. Survival analysis were performed by Mantel-Cox-analysis. In all the cases, the statistical test used is indicated in each figure legends. Results were represented as Mean ± SEM. *p<0.05, **p<0.01, ***p<0.001.

## Results

### Increased gut-liver-translocation in Intestinal loop-experiments in cirrhotic but not pre-hepatic portal hypertensive mice

In control mice as well as in PPVL-animals neither GFP-*E.coli* nor 4 kDa-FITC-dextran did translocate from the ileum to the liver (Fig. 1A). However, 4kDa-FITC-dextran as well as GFP-*E.coli* were detectable in high numbers and hence, significantly increased in translocation to the liver in cirrhotic (BDL and CCl4-treated) mice (Fig. 1B). This was also confirmed in-vivo by applying the dual band laserendomicroscopy to the liver 1 hour after loading the intestinal loop with GFP-*E.coli* (Suppl.Fig.1). In animals with prior ligation of the thoracic duct number of GFP-*E.coli* observed intrahepatically was not significantly altered (Fig. 1C) suggesting that penetrating intestinal bacteria reach the liver mainly via the portalvenous circulation <u>within the protocol applied here.</u>

**Fig. 1:**
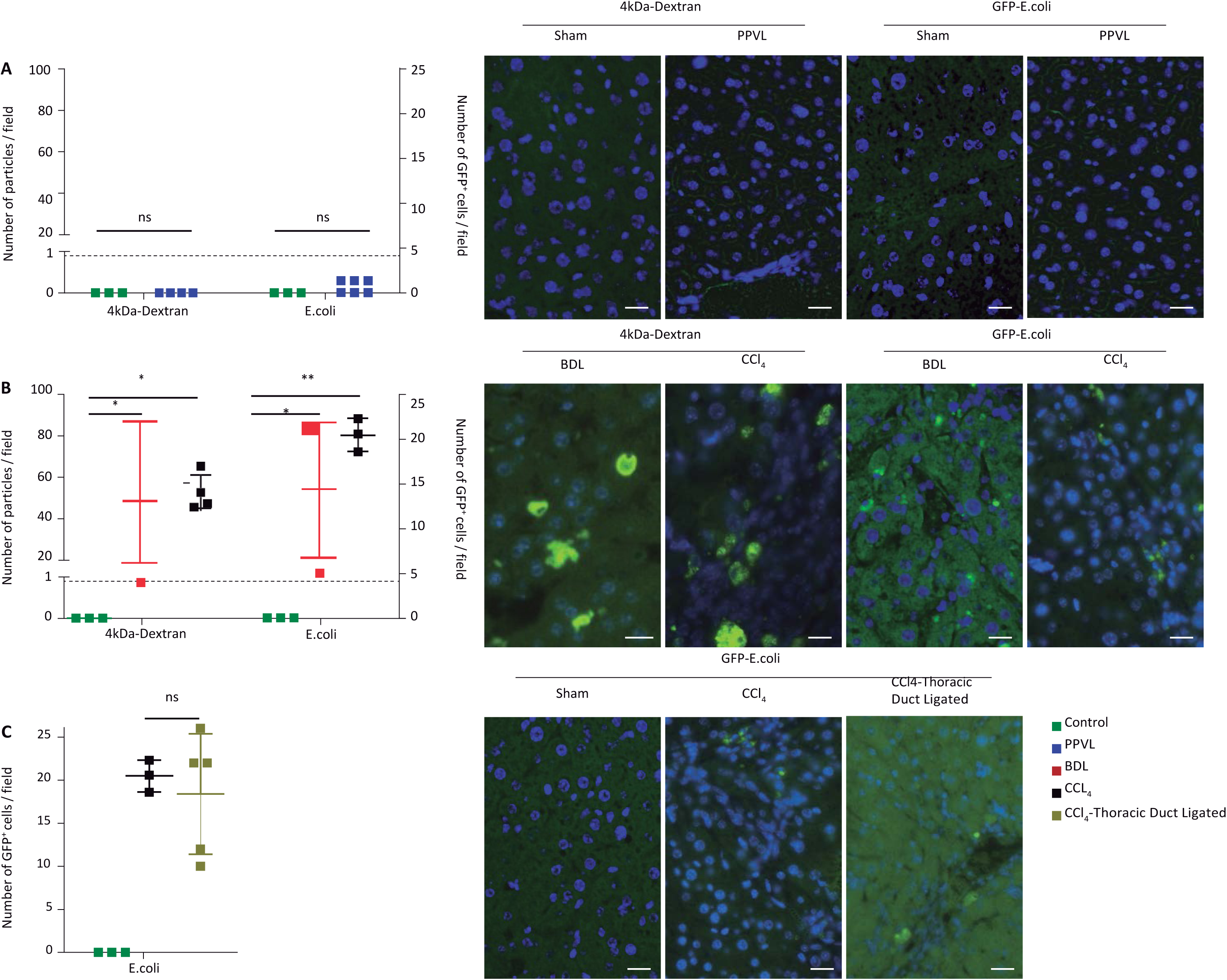
Pathological translocation via gut-liver-axis in cirrhotic but not pre-hepatic portal-hypertensive mice independent from lymphatic route. Liver sections assessed for 4 kDA-FiTC-dextran and GFP-*E.coli* in intestinal loop experiments. A: no detectable fluorescent probe in livers of control or PPVL-mice. B: High retrieval rate of FITC- and GFP-signal in livers of BDL and CCl4-cirrhotic mice being not different in quantity after ligation of the thoracic lymphatic duct (C). Right side corresponding to A-C each representative images. White scale line indicates 20 μm. Colors depicting experimental groups are shown on lower right. <u>Each dot represents the mean counts of at least 10 pictures per animal. Pictures are representative of the indicated experimental groups.</u>

### Lymphatic route of bacterial translocation is physiologically active and increased in intestinal loop-experiments in BDL cirrhotic but not in pre-hepatic portal hypertensive mice

In healthy mice 4kDa-FITC-dextran (not shown) as well as GFP-*E.coli* did translocate from ileal loop to mesenteric lymph nodes (MLN) and peyer patches (PP) demonstrating a physiological low rate of translocation along the lymphatic route (Suppl. Fig. 2A). This lymphatic bacterial translocation was not increased in PPVL mice but augmented in BDL mice (Suppl. Fig. 2A). Interestingly, quantity of GFP-*E.coli* translocation to PP was about 2-fold increased as compared to MLN. Moreover, surprisingly supernatant from PP but not MLN contained high numbers of GFP-*E.coli* indicating cell-independent passage and translocation (Suppl. Fig. 4C). However, severity of translocation to MLN appeared to be less pronounced as compared to gut-liver-axis (Suppl. Fig. 2B). Internalization analysis shows, that cellular fraction of MLN and PP from BDL mice have more GFP internalized than observed in healthy controls and PPVL mice (Suppl. Fig. 4 A,B). The subgroup analysis of GFP-transporting cells revealed the highest frequency among CD11c+/ myeloid dendritic cells (Suppl. Fig.4 D,E). In addition, in cellular fraction of MLN from BDL mice significantly more neutrophils were observed as compared to corresponding evaluation in sham and/or PPVL mice. Analysis of all cellular fraction that doesn’t have GFP internalized (or attached) revealed significantly more neutrophils and CD11c+/myeloid dendritic cells in MLN and PP from BDL mice than from healthy controls and/or PPVL mice (data not shown).

### Mucus layer is reduced in thickness with loss of goblet-cells and MUC2-expression associated with bacterial overgrowth in inner mucus layer in cirrhotic but not PPVL-mice

Considering the increase in goblet cells number in the small intestine from proximal to distal (data not shown) with highest numbers in the terminal ileum we focused on this site for any further investigations. We observed a marked reduction in mucus thickness and goblet cell numbers in germ-free mice (Fig.2A) underlining the role of the microbiota as stimuli for the mucus compartment. Villus morphology in terms of length and perimeter however, was not changed in germ-free conditions (Suppl Fig.5A). Moreover, distribution of goblet cells along the villus with more numbers per length of villus within the base as compared to the apex as well as almost lack of empty appearing GC was similar in germ-free and colonized mice (Suppl Fig.5B, Fig. 2B). In germ-free condition short-term BDL mice (Fig. 2C) presented with reduced mucus thickness. However, <u>BDL mice presented with increased mortality in germ-free conditions (Suppl Fig.6) preventing further investigations</u>. Thus, we utilized a stable moderately diverse mouse microbiota (sDMDM2) as gnotobiotic animal model consisting of pre-defined 12 bacterial species^40^. This microflora also increased GC counts and mucus thickness in terminal ileum of healthy mice enabling a more physiological status. In more detail, this increase in GC in response to this more diverse microflora resulted in an increased number of mucin-filled GC being predominantly pronounced in the base of the villus. Cirrhotic mice under sDMDM2-conditions presented with lower mucus thickness, mucus weight per ileum length as well as goblet cell numbers in terminal ileum as compared to healthy controls (Fig. 2G,H,I, Suppl Fig.5C) with predmoninantly reduction in mucin-filled goblet cells (Fig 2J, Suppl. Fig 5D). Moreover, BDL-cirrhosis resulted in more than 100x times increased bacterial burden in the inner mucus layer (Fig. 2F) evidencing a closing in of bacteria on the epithelial surface normally being almost devoid of bacteria. <u>Also microbial composition changed in BDL-mice presenting with increased abundance of particularly Enterococcocus within the mucus compartment as compared to sham mice (Suppl. Fig 7). On the other hand, decreased abundance of Verrucomicrobia abd Parasutterela was noted in ileal mucus of BDL-mice. Surprisingly, also in pre-hepatic portal hypertension per se (in PPVL-mice) minor dysbiosis was observeable but no change in abundance of enterococci (Suppl. Fig. 7C).</u>

**Fig. 2:**
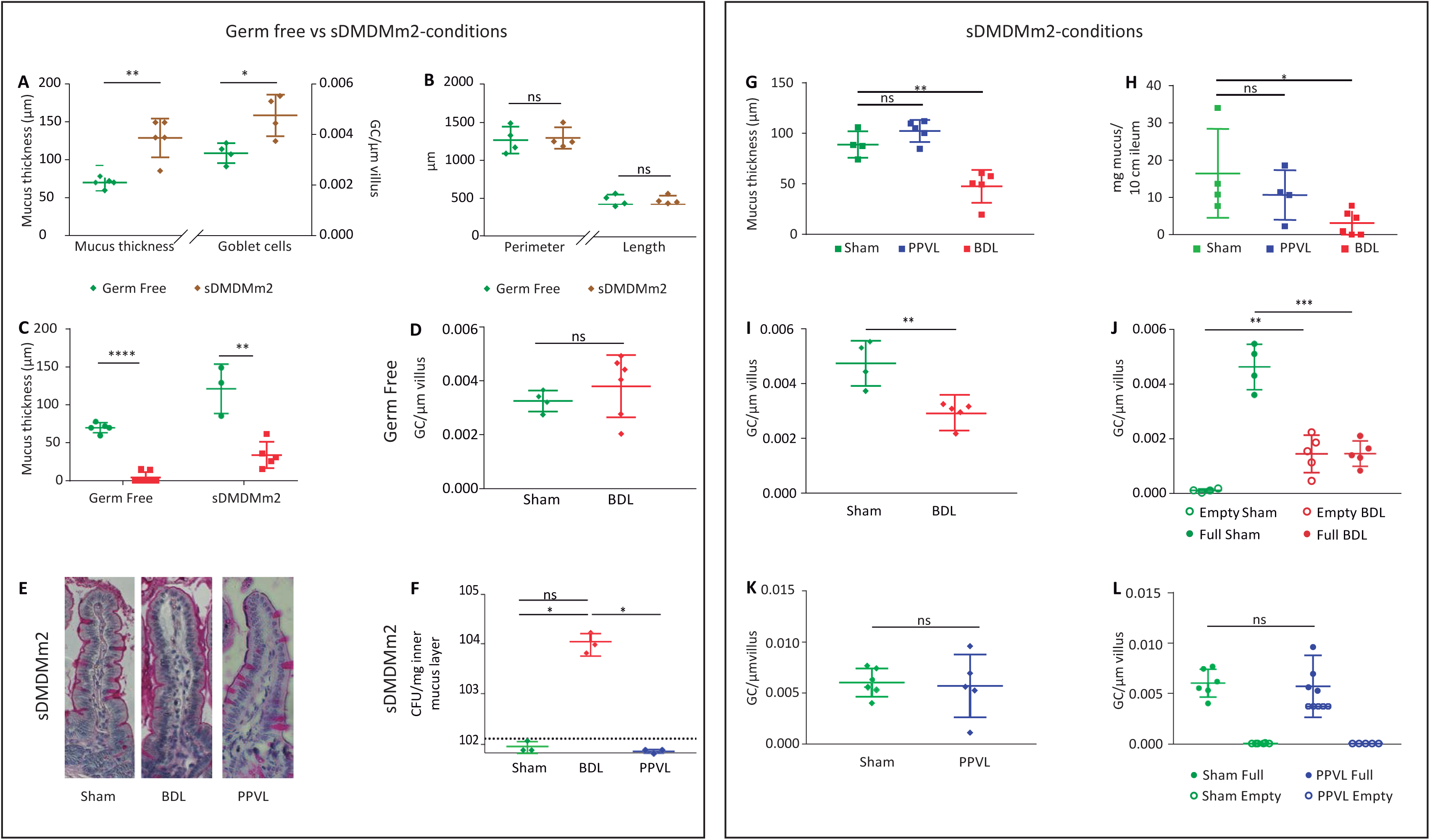
Impact of microbiota and cirrhosis on ileal mucus layer. Left (germ-free vs. sDMDM2). Mucus thickness and goblet cell numbers were reduced in germ-free as compared to sDMDM2-colonized mice (A), and differences in goblet cell numbers related dominantly to those being mucin-filled whereas empty appearing GC were almost absent in germ-free and sDMDM2-conditions (B). In BDL mice already 3 days after surgery mucus thickness was reduced in germ-free as well as in sDMDM2-conditions (C) whereas GC numbers were not affected in germ-free conditions (D) but likewise diminished in sDMDM2-conditions (I). In chronic 14day-PPVL mice in sDMDM2-conditions no change in mucus thickness (G), mucus weight per segment ileum (H) nor GC numbers (total (L) as well as full and/or empty appearing (M)) was noted. In contrast, in BDL mice under sDMDM2-conditions showed increased numbers of empty appearing and reduced mucin-filled GC (J). Representative stainings of ileal villus derived from sham, PPVL and BDL are depicted (E). Inner mucus layer harvested separately after removal of outer layer by suction revealed vastly increased bacterial overgrowth in this normally sterile layer as seen in sham and PPVL mice (F).* p<0.05 and ** p<0.01. t-test; ns: non-significant. <u>Each dot represents one mouse with a mean of at least 30 villi counted. Mean ± standard deviation (SD) displayed for the indicated experimental groups.</u>

In cirrhotic mice, the ones with the lowest mucus thickness within the terminal ileum under sDMDM2-conditions presented with most pronounced bacterial translocation to the liver (Suppl Fig.5E). Contemporarily, cirrhotic animals with pathological translocation to mesenteric lymph nodes or with ascites presented with reduced numbers of goblet cells (Suppl Fig.5 F,G). This indicates that with advanced stage of disease and/or presence of pathological BT goblet cell numbers decreases. Under sDMDM2-conditions total goblet cell count and mucus thickness was not only diminished in long-term and cirrhotic BDL mice but already 3 days after BDL (Fig. 2C) and hence independent of fibrosis/cirrhosis. In that regard, we observed particularly reduced numbers of full but a trend to more empty goblet cells (Fig 2J). In contrast, in germ-free conditions 3 days after BDL no change in goblet cell numbers was noted (Fig. 2D). As for the role of portal hypertension per se no differences in terms of mucus thickness, mucus weight per segment ileum as well as total numbers and mucin-filled or empty goblet cells were observed in PPVL mice as compared to control animals (Fig. 2 G,H,L,M).

RNA-sequencing of whole ileal tissue revealed up-regulation of genes involved in bile acid metabolism, chemotaxis and T-cell recruitment and down-regulation of genes involved in phagocytosis, antigen-presentation, lipid metabolism and goblet cell/intestinal barrier regulation in BDL-mice (Fig. 3A). Noteably, down-regulation of the main mucin-genes (MUC2,-3,-4,-13) was observed suggesting a lack of mucin-synthesis in ileal goblet cells in BDL mice (Fig. 3C). In contrast, no changes in mucin-gene-expression (except for MUC13) could be observed in PPVL-mice (Fig. 3B, C). However, ileal expression of antimicrobial peptides REG3Band REG3G were upregulated in BDL mice without detectable changes in PPVL mice(Fig. 3D).

**Fig. 3:**
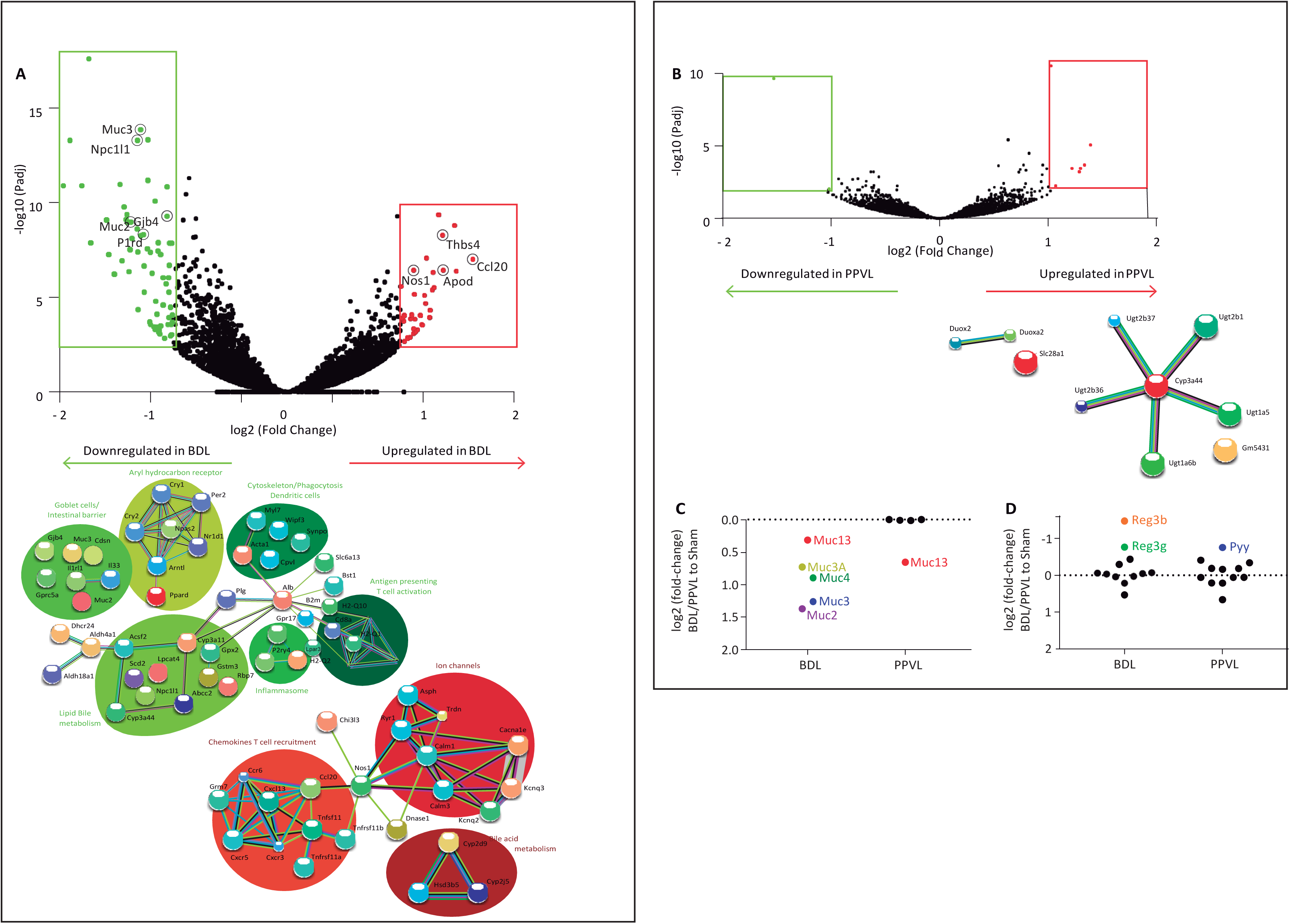
Volcano Plot and String-Analysis of differently expressed genes in terminal ileum tissue. Results show fold-change of selected transcripts between BDL and sham mice (A) and PPVL (B) vs. sham mice. Genes being expressed at significantly altered level are highlighted in red (p-adj<0.01) and genes of interest labeled. String analysis for protein and protein interaction (depicted as connexions) is done on differently expressed genes (log2-fold change>2; p-adj value<0.01) (A and B, lower panel, respectively). Logarithmic Fold-Change in selected transcripts of mucus-synthesis (C) and antimicrobial molecule-(D) expression in BDL and PPVL-mice in relation to sham-animals. Differently expressed genes are colored and labeled (p-adj<0.01). <u>The analysis was conducted using mean gene counts of 3 animals per group.</u>

### Interepithelial leakage and bacterial translocation across the epithelial barrier into the lamina propria are enhanced in cirrhotic but not in pre-hepatic portal hypertensive mice

Dual band confocal laserendomicroscopy in-vivo (Fig. 4) as well as confocal microscopy ex-vivo (data not shown) evidenced translocation of GFP-*E.coli* from the ileal lumen into the lamina propria in BDL-mice. Astonishingly even at high-concentrations of GFP-*E.coli* within the intestinal loops no clear translocation of bacteria across the well-visualized epithelial barrier was noted in healthy control mice during the 30 minute observation period (Fig. 4B). Interepithelial permeation of 70kDA- and 150kDA-FITC-dextran into ileal epithelial layer was not detectable in any control animal (Fig. 5 A,B). In contrast, in cirrhotic (BDL and CCl4-treated) mice but likewise not in PPVL-mice interepithelial leakage for 70 kDa-FITC-dextran was observed (Fig.5A). Correspondingly, PPVL did not show any increase in fecal albumin (66-70 kDA) whereas cirrhotic mice excreted high amounts of albumin demonstrating a disruption of the epithelial barrier (Suppl Fig. 8). Large sized 150kDA-FITC-dextran did not leak into the interepithelial space to the degree defined per protocol to indicate epithelial barrier dysfunction in any of the study groups (Fig. 5B).

**Fig. 4:**
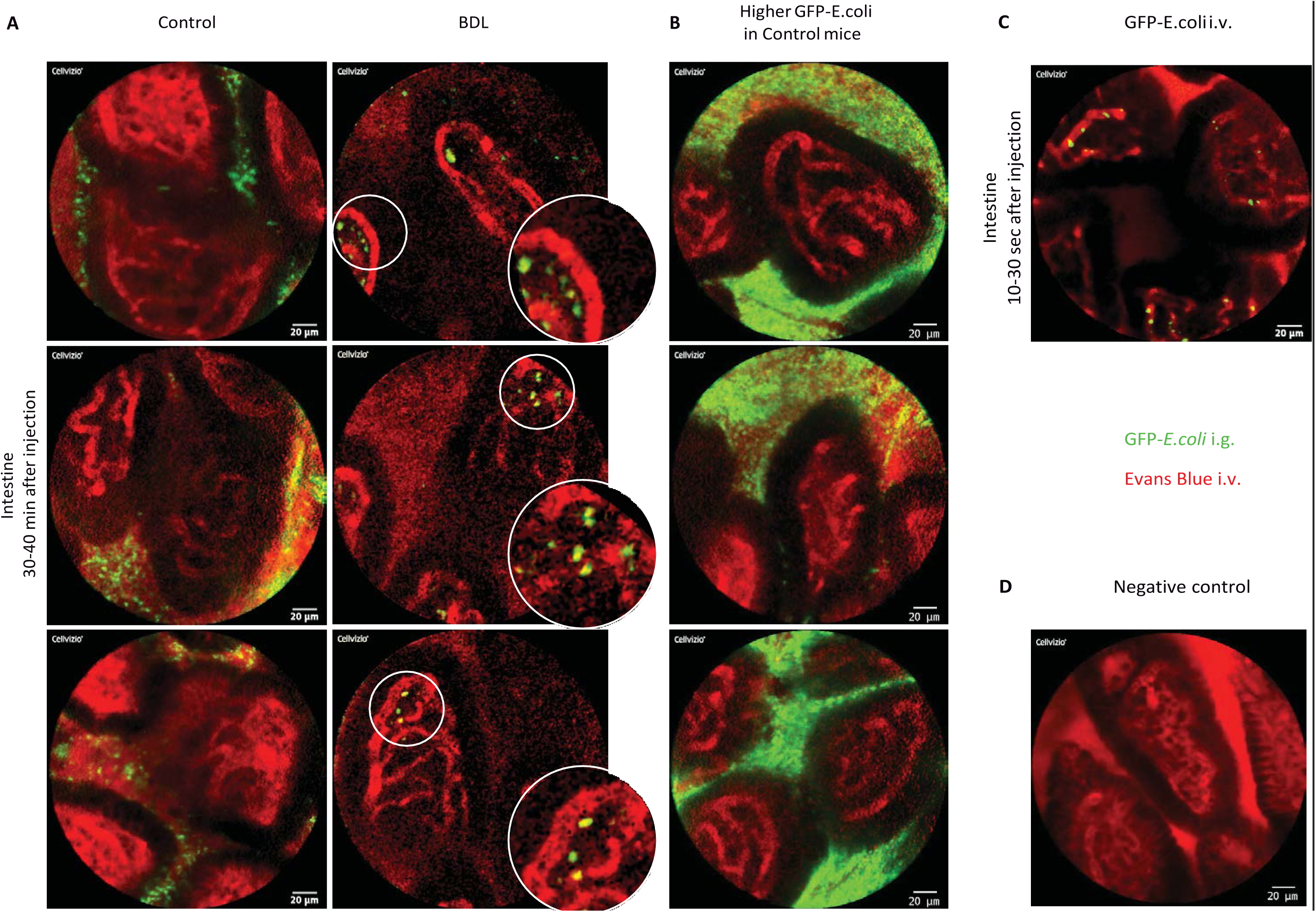
Pathological transepithelial GFP-E.coli-translocation into the lamina propria occurs in BDL-mice but not in healthy conditions or in PPVL mice. Dual-band laserendomicroscopy in-vivo in healthy control and BDL-mice after injection of GFP-*E.coli* into ileal loops intestinal villi were observed for 60 minutes in-vivo showing no GFP-signal in control but multiple translocating GFP-marked *E.coli* within the lamina propria in BDL-mice (A). A sticking of bacteria to the outerside of capillaries can be appreciated with single cases of bacteria having migrated into the vessels is depicted. Even at artificial high concentration of GFP-*E.coli* up to 10^8^ CFU/loop no transepithelial translocation of bacteria into the lamina propria was observed in healthy controls (B). Positive control displaying GFP-*E.coli* within the vasculature after i.v. injection (C) and negative control without exposition to GFP-*E.coli* (D) are shown as well. <u>Pictures are representative of the indicated experimental groups from 2 replicate experiments.</u>

**Fig. 5:**
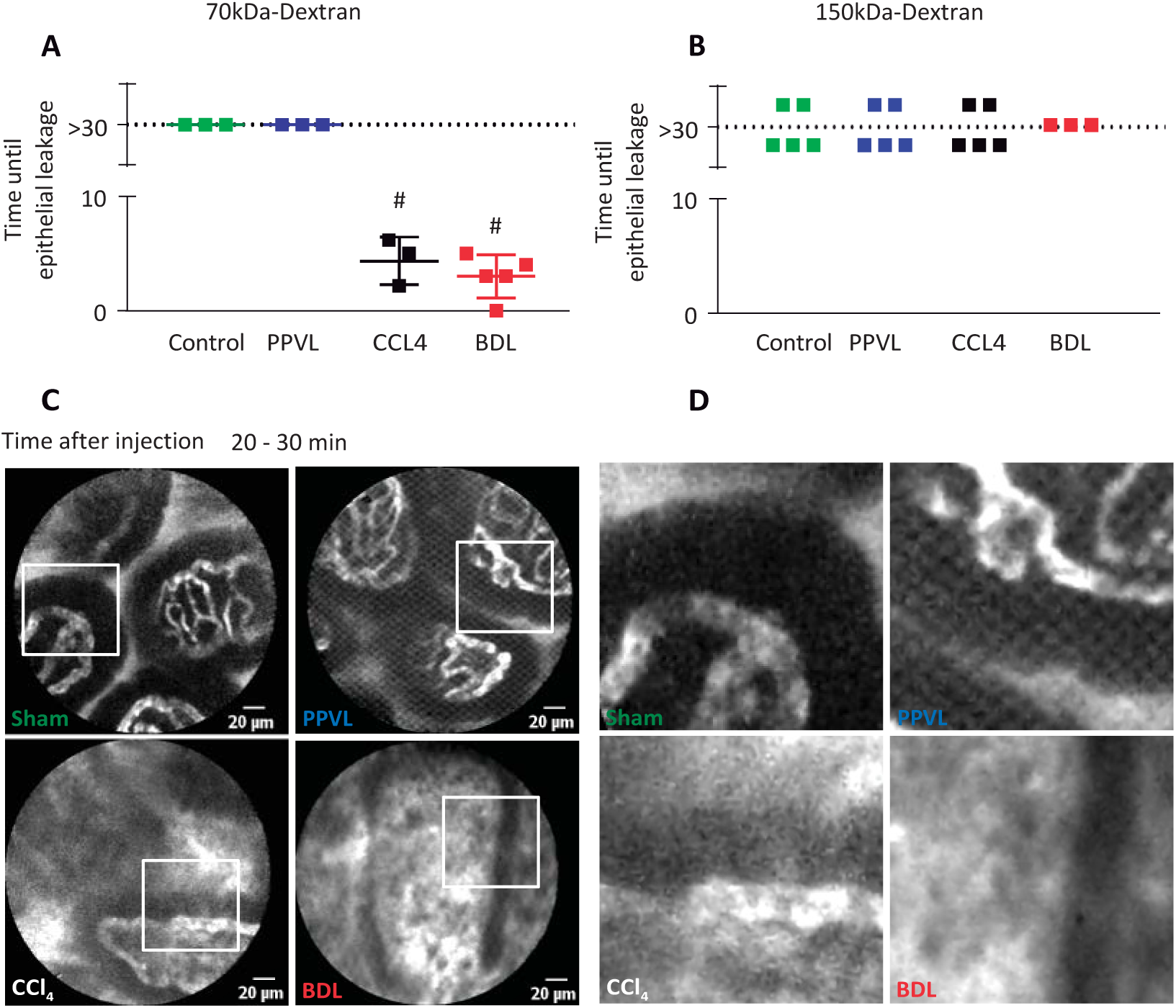
Inter-epithelial leakage of fluorescein is increased in experimental cirrhosis. Pathological interepithelial appearance of 70kDA-FITC-dextran (A) but not 150 kDA-FITC-dextran (B) in ileal villi in cirrhotic BDL and CCL4-treated but not in PPVL-and healthy control mice assessed by pCLE in-vivo. Representative images from confocal laser-endomicroscopy are depicted at the stated observation time points (C) and corresponding magnifications of inlets (D). *** p-value<0.001 T-test. <u>Each dot represents the mean counts of at least 8 pictures per animal and timepoint, the experiment was done at least 3 times. Pictures are representative of the indicated experimental groups and timepoints.</u>

A semiquantitative analysis of IHCs revealed the down-regulation of main tight-junction proteins in ileal epithelium including ZO1, occludin and claudin-1,-2 in CCl4-induced cirrhotic mice compared to naïve control mice (Suppl. Fig. 9). Cirrhotic mice treated with OCA showed a significant increase in all TJ proteins, either reaching or approaching values present in naïve control mice. Cirrhotic mice treated with FEX showed similar results, significantly increasing protein expression of all TJ proteins, except occludin, compared to cirrhotic untreated mice.

### Gut vascular barrier dysfunction in cirrhotic but not pre-hepatic portal-hypertensive mice are independent of intestinal microbiota

In healthy control animals under SPF conditions we could clearly see that 4kDa-FITC-dextran extravasated immediately after injection into the lamina propria (Suppl. Video 1), while 40kDa stayed transiently in the vessel but leaked progressively through the endothelium within the first 3 to 5 minutes reaching a 1:1 ratio after around 10 minutes (Fig. 6B). 70kDa Dextran-FITC could stay in the vessel for up to 8 minutes staining capillaries and giving contrast between vessels and lamina propria, although also extravasated between 5 and 20 minutes. Finally 150kDa-Dextran-FITC remained intravascular for all the observation (30 minutes) in control mice (Fig. 6B). Ileal vascular permeability was increased in cirrhotic (BDL and CCl4) mice with early and enhanced extravasation of 70 kDA (Fig. 6E) and 150 kDa-FITC-Dextran (Fig. 6F) as compared to control mice. In contrast, none of the PPVL-mice did present with augmented leakage of 150-kDa-FITC-dextran being comparable to sham-mice (Fig. 6 F). In germ-free control mice almost identical kinetics and severity of leakage of pre-defined FITC-dextrans as compared to SPF-conditions was observed (Fig. 6A,B). Namely, small-sized 4 kDA- and 40 kDA-FITC-dextrans did permeate early and severly whereas large-sized 150 kDA-FITC-dextran remained intravascularly during the whole observation period. Only 70 kDA-FITC-dextran did extravasate earlier and more severe in germ-free as compared to SPF-mice. Short-term BDL-mice in germ-free conditions presented with vastly disrupted GVB exhibiting rapid and severe extravasation of even large-sized 150 kDA-FITC-dextran (Fig. 6 C,D).

**Fig. 6:**
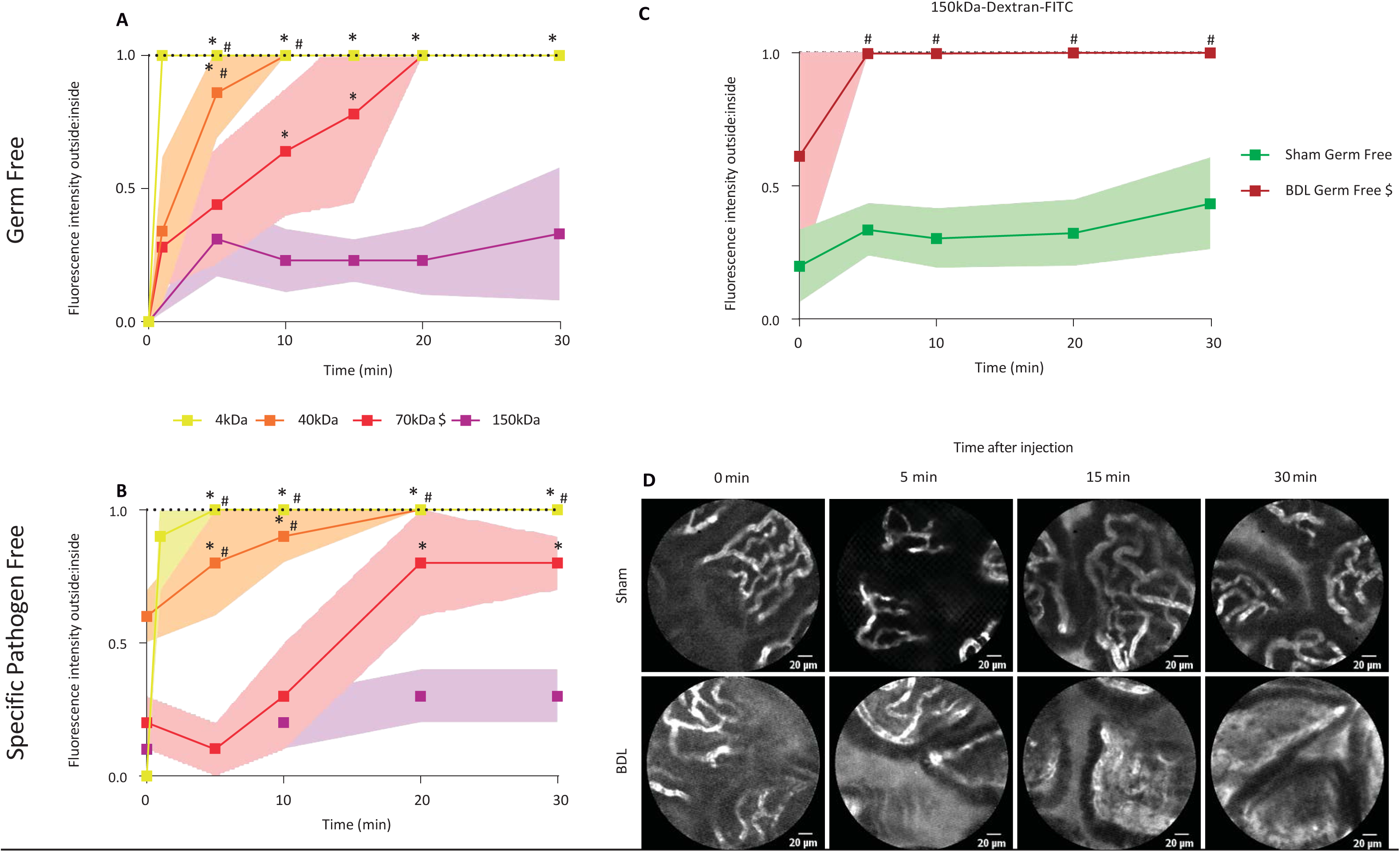
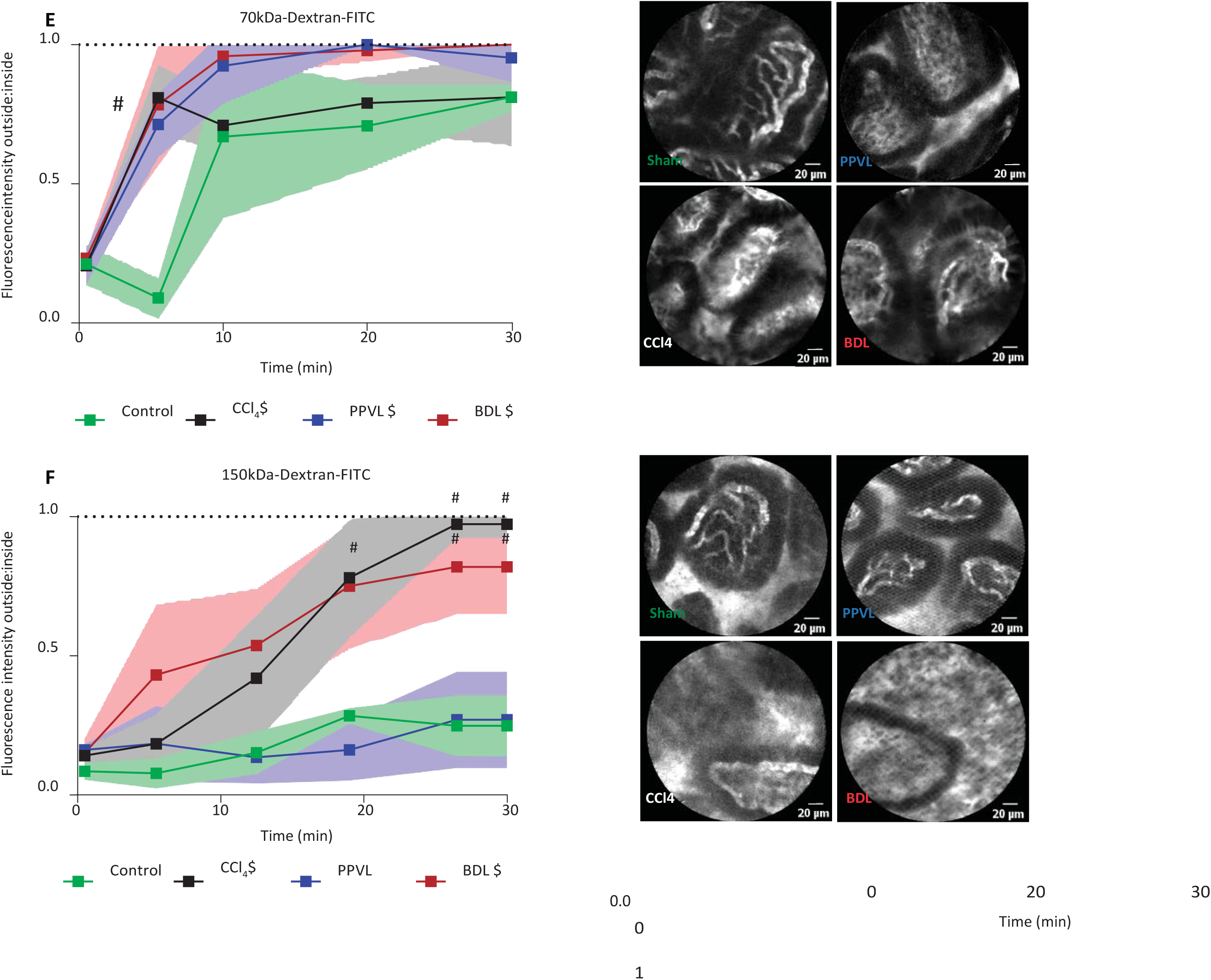
Gut-vascular barrier function in dependency on microbiota and its disruption in cirrhotic mice. **Upper Panel:** Extravasation of different sized FITC-dextran molecules during 30 minutes observation period in healthy germ-free (A) and SPF-mice (B). A very similar kinetic of extravasation can be appreciated for 4, 40 and 150 kDA-FITC-dextran in germ-free and SPF-mice can be appreciated. BDL-mice being born and raised germ-free exhibited grossly accelerated and severe extravasation of 150 kDA-FITC-dextran not being observed at all in sham-mice (C). Representative images at the stated timepoints in germ-free BDL or sham mice (D). **Lower Panel:** Gut-vascular barrier dysfunction for 70-kDA-FITC-dextran (E) and 150-kDA-FITC-dextran (F) in terminal ileum of cirrhotic and pre-hepatic portal hypertensive mice. <u>Statistics: One-way ANOVA corrected by Bonferroni *p-value<0.05 compared to 150kDa; #p-value<0.05 compared to 70kDa. $ Area under the curve significantly different compared to control (t-test: p-value<0.05). Each dot represents the mean counts of at least 8 pictures per animal and timepoint, the experiment was done once. Pictures are representative of the indicated experimental groups and timepoints</u>.

### Increased PV1-expression and changes in endothelial cell morphology correlate with disrupted GVB in cirrhosis

Ileal gut vascular barrier impairment associated with increased PV-1 expression along the gastrointestinal tract. Interestingly, gut vascular analysis in terms of immunofluorescence for PV-1 expression showed increased staining in all small and large bowel analyzed locations in CD34-positive vessels in cirrhotic BDL-mice (Fig.7 A,B). PVL-mice however, displayed minor enhancement in ileum and significanct increases in colon as compared to sham mice (Fig 7 C,D). Electronmicroscopy revealed no major changes in number of fenestrae, caveolae or perimeter of capillaries assessed in cirrhotic or pre-hepatic portal-hypertensive capillaries (Fig. 8A-C). However, morphologically endothelial fenestrae appeared enlarged and inter-cellular membranes reduced in electrodensity in cirrhotic ileal capillaries (Fig. 8D).

**Fig. 7:**
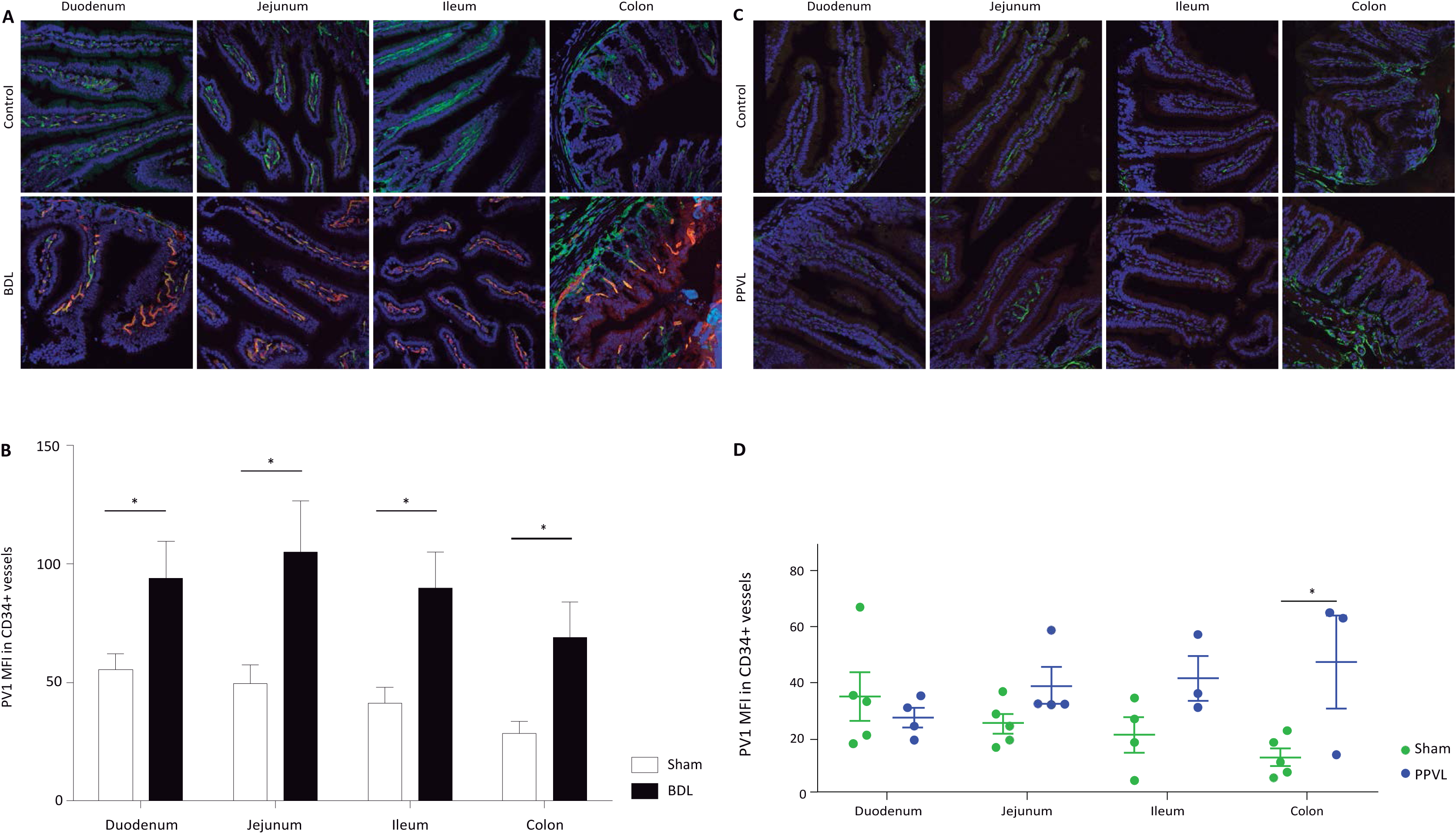
Plasmalemma-vesicle-associated-protein (PV)-1 immunofluorescence in terminal ileum is increased in cirrhotic but only mildly accentuated in PPVL-mice. PV-1 (red) and CD34 (green) staining was performed in different parts of the small intestine (Duodenum, Jejunum, Ileum, and Colon) in BDL (A) and PPVL (C) mice. Quantification of PV-1 fluorescent intensity in CD34 vessels revealed in cirrhotic BDL mice increased staining of PV1 as compared to healthy controls along the small intestine (B). In contrast, in PPVL-mice a minor but consistent increase in PV1 at the ileum and particularly colon was detectable (D). One-way ANOVA corrected by Bonferroni: * p-value<0.05 (A). T-test: ns-non-significant. <u>Each dot represents the mean counts of at least 5 pictures per animal.</u>

**Fig. 8:**
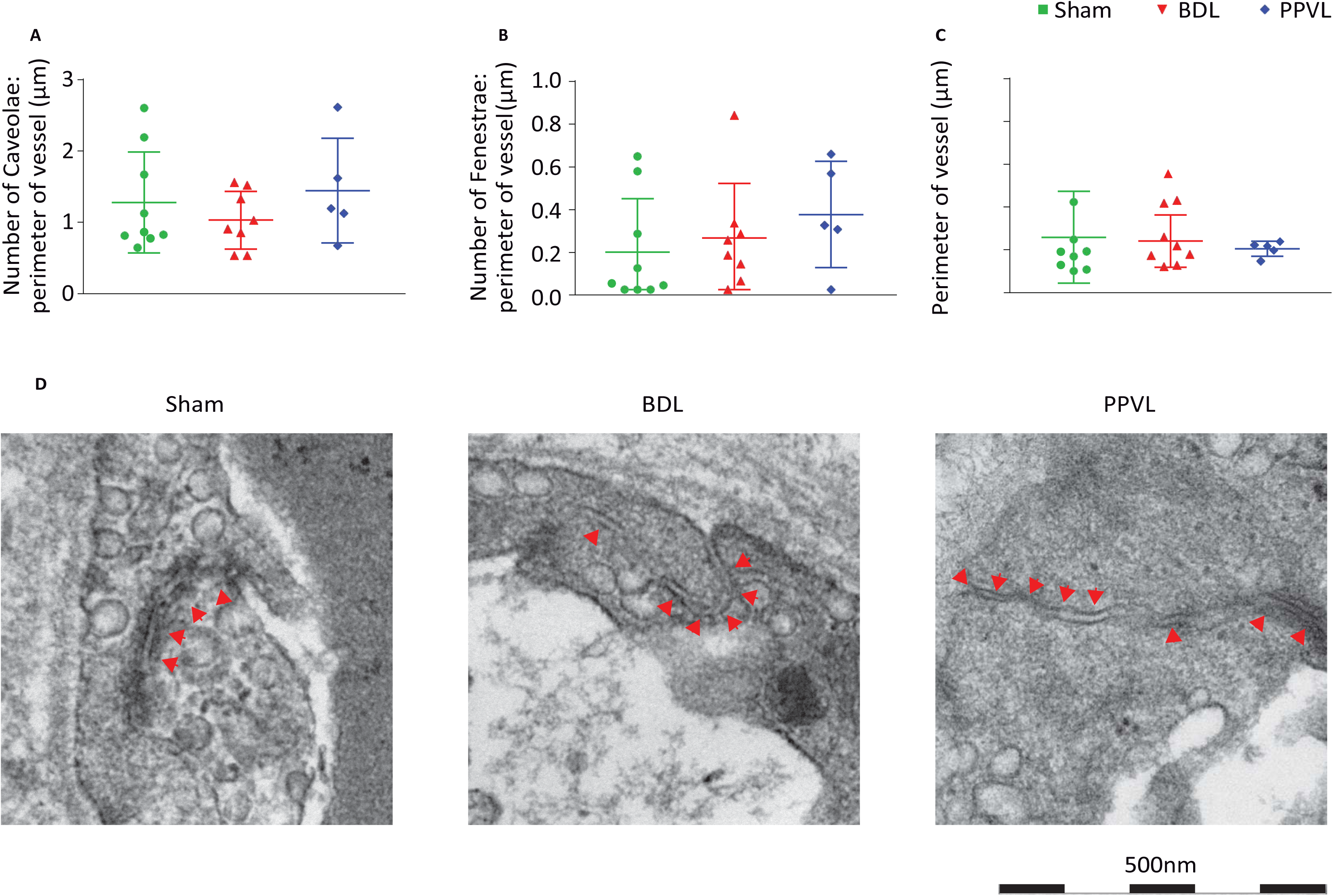
Morphological characteristics of endothelial cells within the ileal microvasculature in BDL cirrhotic and PPVL mice: Number of caveolae (A) and fenestrae (B) and vessel perimeter (C) are measured in ileal villus capillaries in BDL-induced cirrhotic mice, pre-hepatic portal hypertensive mice (PPVL) and sham operated control mice. Each dot or column of dots represent one vessel measured in one mouse respectively (n=2 Sham, n=2 BDL, n=1 PPVL) (A). Representative electron microscopy pictures showing reduced junctional overlapping (red arrow heads) in portal hypertensive mice (D).

### Role of FXR for ileal mucus and gut-vascular barrier and thus pathological translocation along liver-gut-axis

Oral treatment with OCA as well as fexaramine significantly ameliorated pathological translocation of GFP-*E.coli* from the ileal lumen to the liver in cirrhotic mice (Fig. 9 A,B) underlining the impact of pharmacological FxR-activation on “sealing” the access to the gut-liver-axis. However, none of FXR^ΔIEC^ mice analysed did present any translocation of even small 4kDa FITC-dextran or GFP-*E.coli* to the liver (data not shown). In contrast, performing PPVL on FXR^ΔIEC^ mice revealed a small but evident increase in 4kDa FITC-dextran recovery intrahepatically not observed in wild-type PPVL-mice (Fig. 9 C,D). Also in individual PPVL-FXR^ΔIEC^–mice GFP-E.coli was retrievable intrahepatically (data not shown).

**Fig. 9:**
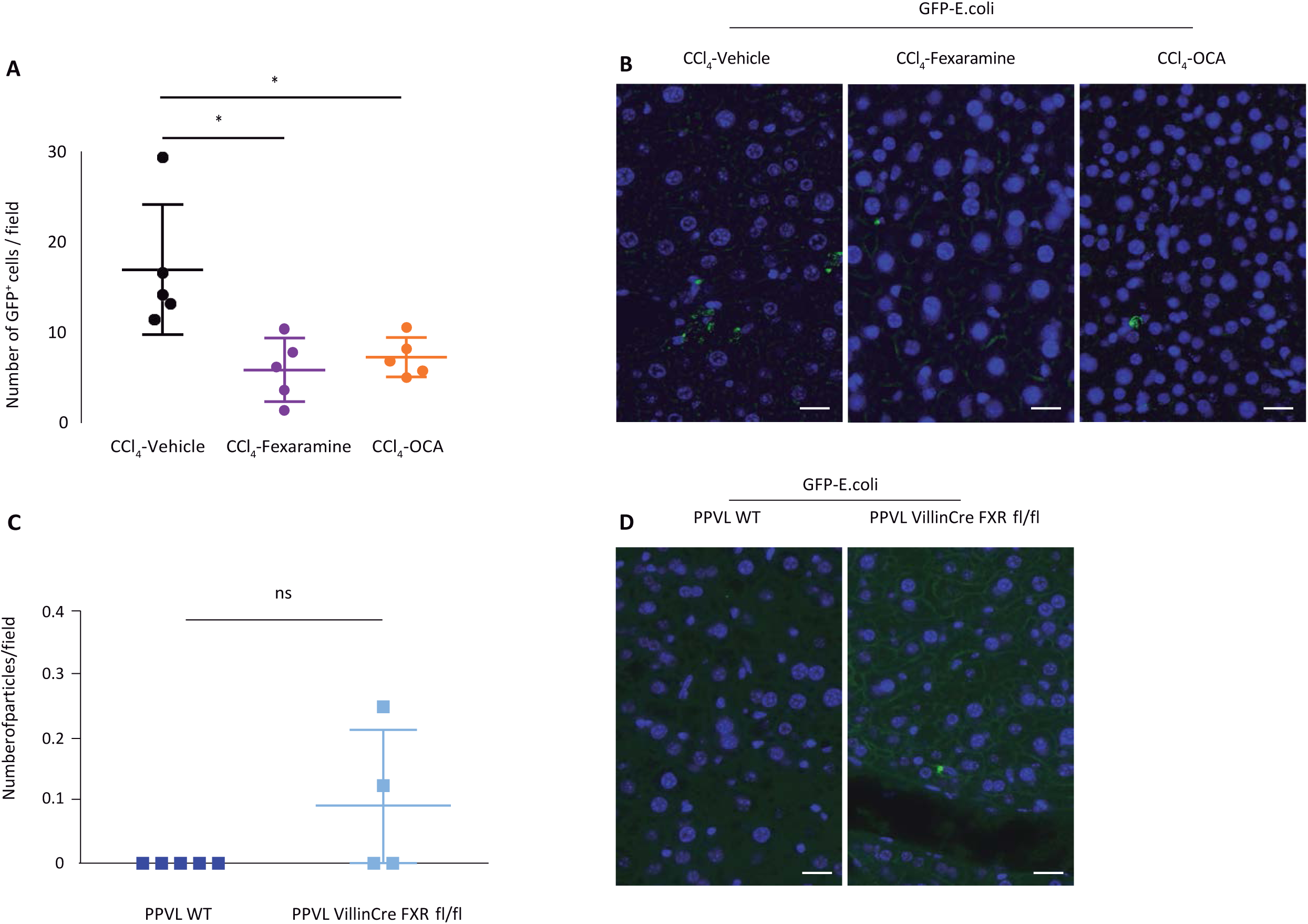
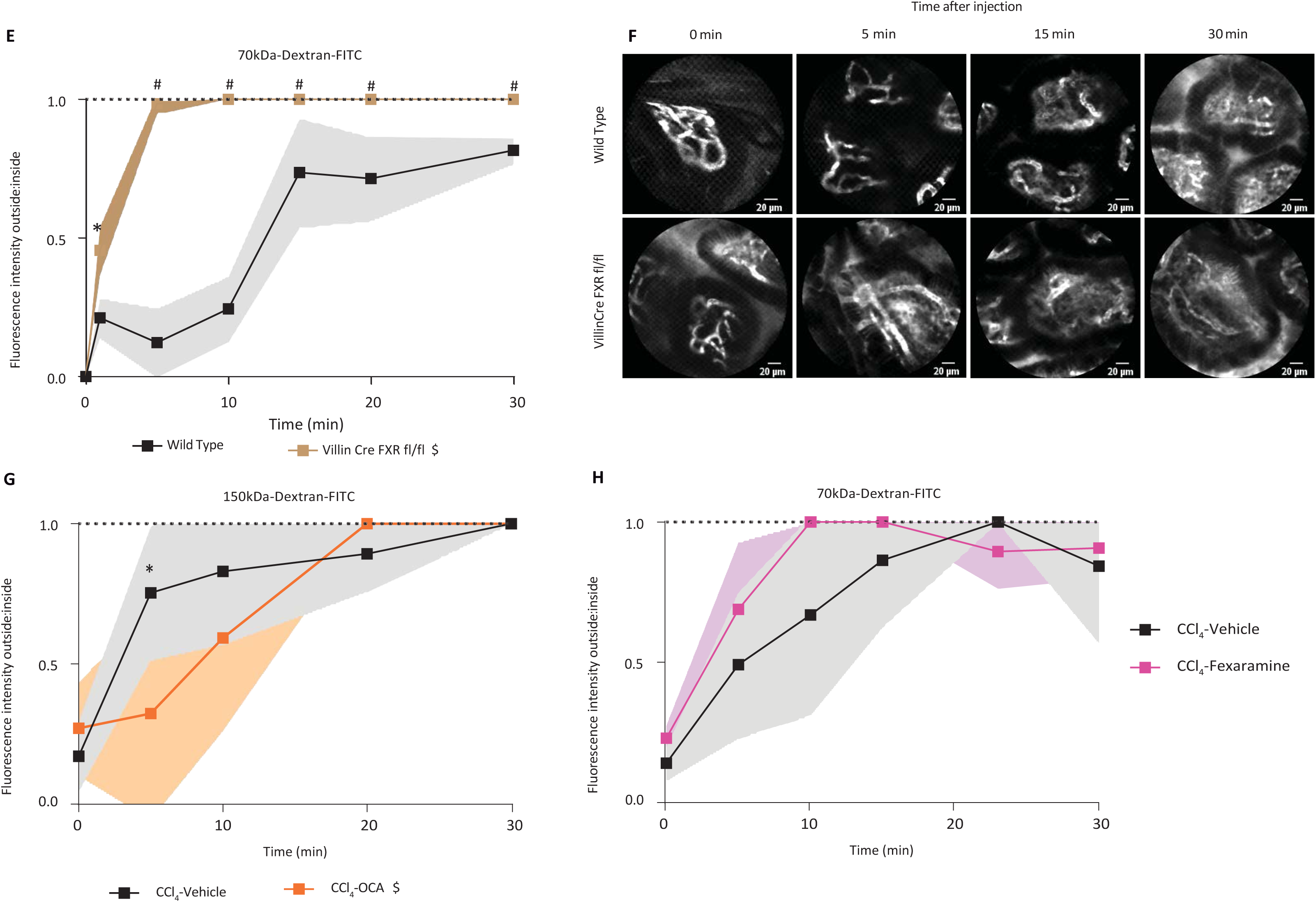
Role of FxXR for gut-liver-axis and gut-vascular barrier. Upper Panel: FXR-agonists fexaramine and obeticholic acid (OCA) reduced hepatic recovery of luminally applied GFP-E.coli (A,B). Quantification of number of bacteria cells in liver sections per field of view 0,5mm^2^. Each dot represents one mouse. Mean ± SD (A, C). Representative pictures of the indicated groups. GFP-*E.coli* (Green) DAPI (blue). White scale lines indicate 20μm (B,D). A,B: CCL4-Induced cirrhotic mice treated for 2 weeks with fexeramine or OCA orally. C,D: PPVL assessed 14days after surgery in wild-type and intestinal-specific FXR-null-mice. One-way ANOVA corrected by Bonferroni: *p-value<0.05. **Lower Panel: Gut-vascular barrier disruption and stabilization in dependency on FXR-singaling. In intestinal-specific FXR-deficient mice GVB is disrupted** as 70 kDA-FITC-dextran extravasation occurs earlier and more severe in mice with epithelial-specific deletion of FXR in the intestine as compared to wild-type-mice (E). OCA-treatment delayed and initially reduced 150-kDA-FITC-dextran extravasation in CCl4-induced cirrhotic mice (G). In contrast, even 70 kDA-FITC-dextran extravasation was not altered by fexaramine-treatment (H). One-way ANOVA corrected by Bonferroni: # p-value<0.001 * p-value<0.05. T-test of AUC comparison: $ p-value<0.05.

Immunohistochemistry for FXR in ileum revealed that FXR was not expressed in VillinCre +/− FXR fl/fl mice (VillinCre FXR fl/fl on figure legends) but it was expressed in epithelial cells in Villin −/− FXR fl/fl (Wild type on figure legends) (Suppl Fig.10). FXR was expressed in all enterocytes but was absent in the nuclei of all PAS-positive mucin-filled goblet cells (Suppl Fig. 10). Interestingly we also observed a gradual reduction of FXR expression from the apex of the villus where the expression was higher to the base of the crypt. This absence of FXR expression is specifically more pronounced at the base of the crypt in which stem cells and Paneth cells are located and the highest concentration of goblet cells is found.

As for the muco-epithelial barrier OCA-treatment partly restored goblet cell numbers in cirrhotic animals being no more statistically significant different from control animals (Suppl Fig. 10). In addition, OCA significantly increased ileal tight-junction-protein-expression (ZO1, claudin-1,-2 and occludin) up-regulating them to the level of healthy control mice (Suppl. Fig.9). Fexaramine with almost exclusive muco-epithelial active concentrations likewise did improve tight-junction protein immunohistochemical staining (Suppl. Fig. 9) but failed to improve gut-vascular barrier when challenged with 70 kDA-FITC-dextran (Fig. 9H) in CCl4-mice. In contrast, endothelial FxR-stimulation correspondingly, delivered by OCA as systemic FXR-agonist attenuated, at least partly, GVB dysfunction in cirrhotic mice (Fig. 9G).

## Discussion

In this paper we report changes in mucus and gut-vascular barrier in relation to microbial modulation and as entry sites for pathological bacterial translocation along the gut-liver-axis in cirrhosis. Standardized in-vivo intestinal loop-experiments are utilized to quantify the translocation process from the intestinal lumen to the liver. Pathological increases in bacterial translocation are evidenced in cirrhotic mice occuring largely independent from the lymphatic route. This does not exclude the well-known PBT along the lymphatic route in cirrhosis as demonstrated by us^14, 41^ and multiple other independent investigators^42–44^. <u>We here confirm this in BDL mice and extend this observation by demonstrating a particularly pronounced PBT into peyer patches which appears to involve mainly myeloid dendritic cells but also represents at least partly a cell-independent passage and translocation process.</u> However, translocation into the lymphatic system is a physiological event occurring in healthy conditions via a highly orchestrated process at low rates. In contrast, here we show that no bacterial and/or 4kDA-dextran-translocation is detectable intrahepatically in control and pre-hepatic portal hypertensive mice. This strongly argues for a tight sealing and protection of the gut-liver-axis from any bacterial translocation in healthy conditions which is not significantly altered by portal hypertension. In liver cirrhosis however, with pathological bacterial translocation the gut-liver-axis is markedly “opened” and fueled with bacteria via the intestinal microcirculation and portalvenous route which seems to represent the predominant route of access to the host.

We chose the distal ileum as main target site for investigations due to the most drastic increase of bacterial load and the presence of thinner and more penetrable mucus layer compared to the large intestine^45^. Nonetheless, also in the ileum the mucus layer represents the first line of defense against bacterial translocation with mucus release keeping bacteria at distance from the epithelium^13^. Correspondingly defects in mucus barrier function result in increased bacterial adhesion to the surface epithelium and increased intestinal mucosal permeability^46, 47^. Colonization of the gastrointestinal tract with microbial commensals has been reported before to impact on intestinal mucus machinery as well as vascular development. It has many crucial roles including on normal development of mucus barrier^31^, on induction of mucus secretion, increase the number of goblet cells via activation of toll like receptors in the intestinal epithelium by microbial derived products^48^. Goblet cells have been studied in germ free conditions before and found to be reduced in numbers and smaller in size in cecum as compared to conventionally raised mice^49^. Here we extend this observation for another site namely small intestine and sDMDM2-gnotobiotic conditions demonstrating increased total numbers of goblet cells and particularly those being mucin-filled as compared to germ-free mice. Moreover, several members of the microbiota community have been reported to increase *muc2* gene expression *in vivo* and *in vitro* and to release the otherwise membrane-anchored MUC2 molecule^50^. Here we demonstrate under sDMDM2-gnotobiotic contditions reduced mucus thickness and expression level of main mucus-genes in cirrhotic mice. This associates with bacterial overgrowth in the inner mucus layer elsewise normally being sterile promoting pathological crossing of *E.coli* into the lamina propria. Most translocation of living bacteria to the liver is seen in mice with low mucus thickness and cirrhotic mice with translocation to the mesenteric lymph nodes present with a reduction in goblet cell numbers as compared to those without bacterial translocation. Thus, changes in mucus parameters at least partly can be explained by the observed reduction in goblet cells with a particularly pronounced diminished number of mucin-filled goblet cells and concomitant increase in empty goblet cells in liver cirrhosis under standardized gnotobiotic conditions. Surprisingly, lack of luminal intestinal bile induced by short-term BDL in absence of microbiota failed to impact on goblet cell density. This hints at the microbiome being responsible for the observed changes in sDMDM2-conditions. <u>Thus, we analysed ileal microbial composition in mucus demonstrating changes in microbial clustering with overgrowth of particularly enterococcus faecalis in BDL</u> mice. Considering the mucus-binding as well as mucus degradation capacity of enterococci^51, 52^ <u>it is tempting to speculate that this drastic increase in enterococci might actually contribute to the observed reduction in mucus thickness in these animals. More interestingly, these data shed new light on most recent demonstration of</u> enterococcus overgrowth in gastric acid depleted conditions and associated progression of alcoholic liver disease^53^. Indeed, in conjunction with the observed protection of muc2-deficient mice from alcoholic liver disease^54^ makes the mucus niche a highly attractive target for preventive and/or therapeutic modulations in liver diseases. <u>However, further studies need to address this as well as the exact role of bile acids for the observed changes.</u>

Microbial commensals of the gastrointestinal tract have been implicated in the development of the intricate network of blood capillaries in intestinal villi at least partly depending on angiogenic factors secreted by Paneth cells^32^. More interestingly monocolonization with *Bacteroides thetaiotaomicron* a common human and mouse gut commensal is enough to promote vascular development. Consistently with these results we have also shown that transient bacterial colonization with the reversible auxotrophic *E.coli* strain HA107 is enough to induce angiogenic genes and increase vascular density within the intestinal villi^55^. As for vascular barrier function here we demonstrate that the microbiome is of rather limited impact. In fact, germ-free mice present with similar permeation characteristics for small and large-sized molecules as do mice with normal gut flora. However, a modulatory role for microbes and microbial products cannot be excluded considering the observed expedited leakage of pathological sized molecules in BDL-mice in germ-free conditions. This indeed hints towards some protective effects of the microbiome in stabilizing the gut-vascular-barrier. In fact, also for 70 kDA-dextrans resembling albumin in size extravasation is slower in presence of a complex microbiome as compared to germ-free conditions. Thus the microbiome appears to be of major relevance for the mucus machinery but although exerting some impact gut-vascular barrier appears rather unaffected in permeability.

Vascular changes in the splanchnic circulation have been characterized intensively in terms of compliance, hemodynamic dysregulation and angiogenesis but only limited data are available as for endothelial permeability. The newly described gut-vascular barrier is grossly disrupted in cirrhotic mice (BDL, CCl4) with rapid extravasation of large-sized molecules (150kDA) being more than twice in size than albumin. Concomitantly, enteral loss of albumin is observed in cirrhotic conditions but not in pre-hepatic portal hypertension. This is in accordance with the lack of inter-epithelial fluorescence being detectable for 70kDA-FITC-dextran in PVL-mice considering albumin being 66 kDA in size and known to leak predominantly para-cellularly. The expedited extravasation of large-sized molecules from intestinal capillaries in cirrhotic mice easily explains the well-known edema within the lamina propria due to concomitant fluid shifts and increases in interstitial pressures. Vascular hyperpermeability in cirrhotic mice did associate with augmented PV-1 availability known as biomarker for the level of endothelial permeability^56^. This is in line with our previous observation of increased expression of PV1 and intestinal endothelial permeability during high-fat-diet and evidenced disruption of the gut-vascular-barrier^57^. The exact mechanisms upregulating PV1-availability in cirrhosis need to be delineated but could involve endothelial growth factor-A (VEGF). Indeed, PV1 has been reported to be upregulated by VEGF^58^ and inhibition of PV1-expression resulted in decreased VEGF-induced endothelial permeability of fluorescent tracers, both in vivo and in vitro^59^. Considering the well-known increased serum levels of VEGF in experimental cirrhosis it is tempting to speculate that this contributes to the observed PV1-upregulation.

As for the underlying mechanism of pathological bacterial translocation to the liver we propose that the muco-epithelial and vascular endothelial barrier need to be disrupted simultaneously in order to allow permeation from the lumen into the portal-venous circulation such as in liver cirrhosis but not in pre-hepatic portal hypertension. Indeed, portal hypertension per se in absence of liver injury fails to induce alterations in mucus thickness, -weight per ileum length and MUC2-expression as well as goblet cell numbers. In addition, increased portal pressure alone does not associate with permeation of *E.coli* trans-mucoepithelial into the lamina propria. Moreover, although gut-vascular barrier is disrupted to some degree this is limited to small-/medium-sized molecules up to 70kDA- but not large-sized 150 kDA-dextrans. However, this minor alteration is sufficient to enable some access to the villus microcirculation for molecules normally not permeating the vascular endothelium when a second hit such as lack of epithelial FXR-signaling known to affect epithelial barrier is added. Indeed, FXR^ΔIEC^ mice per se do not present with either 4kDA-FITC-dextran or GFP-*E.coli* translocation to the liver but once PPVL is performed and GVB is disrupted the gut-liver-axis is fueled and 4kDA-FITC as well as *E.coli* do appear intrahepatically.

Thus, FXR appears to play a central role for modulating both barriers explaining the therapeutic benefit of systemic and/or local intestinal FXR-activation established here in experimental cirrhosis. Both FXR-agonists, OCA and fexaramine abrogated pathological *E.coli* translocation to the liver underlining their effect in “sealing” the gut-liver-axis. FXR-agonists have been used before in experimental portal hypertension^15, 43, 60, 61^. However, direct assessment as for their effectiveness on the gut-liver-axis is lacking. OCA has been reported to lower intrahepatic resistance ameliorating portal hypertension^60, 61^ and FXR-activation in the small intestine reduces bacterial translocation to mesenteric lymph nodes in experimental cirrhosis^15, 43^ or acute cholestasis^62^. However, the mode of action in our experiments as for the gut-liver-axis appears independent from portal hypertension and not influenced by the lymphatic drainage. Hence, overall we present a new modulatory role of FXR for mucus and gut-vascular barrier affecting the gut-liver-axis.

FXR activation by OCA as well as fexeramine increased ileal protein expression of main TJ-proteins (ZO1, occluding, claudin-1, -5) in cirrhotic mice. This is in line with previous reports demonstrating FxR-activation to influence intestinal epithelial cell proliferation and apoptosis^25^, increase ileal ZO1-expression^15^ and to exert potent anti-inflammatory actions in the intestine, stabilizing epithelial integrity^26, 63, 64^. Here we report, in addition, that OCA impacted on the mucus-machinery by increasing ileal goblet cell numbers in cirrhotic animals to a level similar to control mice. Moreover, the fact that already 3 days after BDL in absence of liver fibrosis/cirrhosis mucus thickness is reduced and numbers of goblet cells are decreased supports a role of bile salts for goblet cell homeostasis. In conjunction, with the well-known deficiency in luminal availability of bile/acids^24^ and associated reduction in ileal FXR-signalling in cirrhotic rodents^15^, it is tempting to speculate that deficient bile-acid induced FXR-signalling could mediate this phenotype. However, we were unable to detect FXR-expression in goblet cells suggesting a paracrine mode of action for bile/acids which needs to be delineated in further studies.

On the vascular site FXR has been shown to be expressed on endothelial cells^65^ and we have reported previously that specific FXR deletion in endothelial cells disrupts the GVB, leading to liver damage in high-fat-diet-models^57^. Here we extend these data by demonstrating a stabilization of the dysfunctional gut-vascular-barrier in cirrhotic animals by systemic active FxR-agonist OCA. In contrast, Fexeramine failed to impact on the gut-vascular barrier. The latter indicates that improvements in each barrier separately can be sufficient to block pathological bacterial translocation to the liver.

In summary, the microbiota appears key for mucus- and goblet cell physiology whereas its impact on the gut-vascular barrier function seems rather limited. Access to the gut-liver-axis in healthy conditions with normal gut flora is highly regulated and restricted in size and quality. In contrast, pathological translocation to the liver develops in cirrhosis independently from portal hypertension as well as the lymphatic route. This opening of the gut-liver-axis associates with mucus- and GVB dysfunction modulated at least partly by FXR. Obeticholic-acid and fexaramin both reduce pathological bacterial translocation in cirrhosis but acting differently at the muco-epithelial and the vascular endothelial barrier.

## Author contributions

M.S. ideated and performed the experiments; M.J., D.S., Y.N., S.M., M.H. helped M.S. in the execution of the mouse experiments; B.Y. and L.H. performed microbial analysis; M.R. provided human investigations and administered the informed consents; A.G., M.R., A.A. participated with ideas, results interpretation, and careful reading of the manuscript; R.W. ideated the study, coordinated the work, and wrote the manuscript.

## Acknowledgements

The intestine-specific FXR-null (FXRΔ^IE^) mice were a kind gift of Prof. B. Schnabl (University of San Diego, CA, USA); Electron microscopy sample preparation and imaging were performed with devices supported by the Microscopy Imaging Center (MIC) of the University of Bern. We greatly appreciate technical support by F. Blank and C. Wotzkow from Department Pneumology and Department for Biomedical Research, University of Bern, Bern, Switzerland

